# CCA1 and ATAF2 differentially suppress cytochrome P450-mediated brassinosteroid inactivation in *Arabidopsis*

**DOI:** 10.1101/457200

**Authors:** Hao Peng, Michael M. Neff

**Author notes:** Correspondence; Hao Peng.

## Abstract

Brassinosteroids (BRs) are a group of steroid hormones regulating plant growth and development. Since BRs do not undergo transport among plant tissues, their metabolism is tightly regulated by transcription factors (TFs) and feedback loops. BAS1 (CYP734A1, formerly CYP72B1) and SOB7 (CYP72C1) are two BR-inactivating cytochrome P450s identified in *Arabidopsis thaliana*. We previously found that a TF ATAF2 (ANAC081) suppresses *BAS1* and *SOB7* expression by binding to the Evening Element (EE) and CCA1-binding sites (CBS) on their promoters. Both EE and CBS are known binding targets of the core circadian clock regulatory protein CCA1. Here, we confirm that CCA1 binds the EE and CBS motifs on *BAS1* and *SOB7* promoters, respectively. Elevated accumulations of *BAS1* and *SOB7* transcripts in the *CCA1* null mutant *cca1-1* indicate that CCA1 is a repressor of their expression. When compared to either *cca1-1* or the *ATAF2* null mutant *ataf2-2*, the *cca1-1 ataf2-2* double mutant shows higher *SOB7* transcript accumulations and stronger BR-insensitive phenotype of hypocotyl elongation in white light. CCA1 interacts with ATAF2 at both DNA-protein and protein-protein levels. *ATAF2*, *BAS1* and *SOB7* are all circadian-regulated with distinct expression patterns. These results demonstrate that CCA1 and ATAF2 differentially suppress *BAS1*- and *SOB7*-mediated BR inactivation.

**Highlight:** The core circadian regulator CCA1 is a direct repressor of brassinosteroid inactivating genes *BAS1* and *SOB7*, and interact with another repressor, ATAF2. Their differential suppressing effects are regulated by light.

**Abbreviations:** 3-aminotriazole (3-AT), brassinolide (BL), brassinosteroid (BR), CCA1-binding site (CBS), cytochrome P450 (P450), Evening Element (EE), transcription factor (TF), yeast one-hybrid (Y1H), yeast two-hybrid (Y2H)

## Introduction

Brassinosteroids (BRs) are a class of polyhydroxysteroid hormones that regulate plant growth (Zhu *et al*., 2013a), stress tolerance (Nolan *et al*., 2017), and disease resistance (Belkhadir *et al*., 2012). Unlike other phytohormones, BRs do not undergo transport process within the plant body (Symons and Reid, 2004; Savaldi-Goldstein *et al*., 2007; Symons *et al*., 2008). Therefore, their biosynthesis and catabolism are tightly regulated in different plant tissues and developmental stages (Zhao and Li, 2012). In the model plant *Arabidopsis thaliana*, several transcription factors (TFs) have been identified as regulators of the BR biosynthetic genes *DWF4*, *CPD* and *BR6ox2*. A TCP-family TF TCP1 (Guo *et al*., 2010) and a NAC-family TF JUB1 (Shahnejat-Bushehri *et al*., 2016) activate and suppress the expression of *DWF4*, respectively. COG1, a Dof-type TF, binds to the promoters of two phytochrome-interacting-factor-encoding genes *PIF4* and *PIF5* to promote their expression (Wei *et al*., 2017). PIF4 and PIF5 are two bHLH TFs that directly promote the expression of *DWF4* and *BR6ox2* (Wei *et al*., 2017). Two homologous TFs CESTA and BEE1 interact with each other and promote the expression of *CPD* by directly binding a G-box motif in its promoter (Poppenberger *et al*., 2011).

In *Arabidopsis*, the transcription of key BR biosynthetic and catabolic genes is feedback regulated to maintain hormone homeostasis (Tanaka *et al*., 2005). BAS1 (CYP734A1, formerly CYP72B1) and SOB7 (CYP72C1) are two BR-inactivating cytochrome P450s (P450s) that are subject to transcriptional feedback regulation loops (Neff *et al*., 1999; Turk *et al*., 2003; 2005; Thornton *et al*., 2010). Overexpression of *BAS1*, *SOB7* or their orthologs from other plant species confer a BR-deficient dwarf phenotype in *Arabidopsis* (Neff *et al*., 1999; Turk *et al*., 2005; Thornton *et al*., 2011).

Three TFs are known to be the transcriptional regulators of *BAS1* or *SOB7*. LOB directly binds the promoter of *BAS1* and activates its expression (Bell *et al*., 2012). The auxin response factor 7 (ARF7) can bind to the E-box motifs of the *BAS1* promoter and suppress its expression (Youn *et al*., 2016). We previously reported that the NAC TF ATAF2 (ANAC081) can bind to the promoters of both *BAS1* and *SOB7* as a repressor (Peng *et al*., 2015). ATAF2 is also known to regulate disease resistance (Delessert *et al*., 2005; Wang *et al*., 2009b; Wang and Culver, 2012b), abiotic stress tolerance (Takasaki *et al*., 2015), and auxin biosynthesis (Huh *et al*., 2012). ATAF2 can act as either an activator or repressor depending on growth conditions or promoter context (Delessert *et al*., 2005; Wang *et al*., 2009b; Nagahage *et al*., 2018).

ATAF2 binds the Evening Element (EE; AAAATATCT or its reverse complement sequence) and the CCA1-binding site (CBS; AAAAATCT or its reverse complement sequence) on *BAS1* and *SOB7* promoters (Peng *et al*., 2015). The EE sequence has one extra “T” when compared to that of the CBS, and both are known as the binding targets of the core circadian clock regulatory protein CCA1 (Wang and Tobin, 1998; Michael and McClung, 2002; Harmer and Kay, 2005). CCA1 is a MYB TF initially identified as an activator of *Lhcb1*3*, which encodes a light-harvesting chlorophyll a/b protein (Wang *et al*., 1997). Similar to ATAF2, CCA1 can act as either an activator (Fujiwara *et al*., 2008) or a repressor (Li *et al*., 2011) of downstream genes under different circumstances.

In this research, we confirmed that CCA1 binds the EE and CBS elements of *BAS1* and *SOB7* promoters, respectively. Like ATAF2, CCA1 is also a repressor of *BAS1* and *SOB7* expression. CCA1 interacts with ATAF2 at both DNA-protein and protein-protein levels. The suppressing effect of CCA1 and ATAF2 on *SOB7* expression can be either additive or redundant depending on the light or dark growth conditions for *Arabidopsis* seedlings. *ATAF2*, *BAS1* and *SOB7* are all circadian-regulated with distinct expression patterns. Our findings provide novel insight into the connection between BR homeostasis and circadian clock regulatory pathways.

## Materials and Methods

### Plant materials and growth conditions

All *Arabidopsis* plants used in this study are in the Columbia (Col-0) background. The *cca1-1* in Col-0 background (CS67781) and *ataf2-2* (SALK_015750) mutants were obtained from the *Arabidopsis* Biological Resource Center (ABRC). The *cca1-1* in Col-0 was created by backcrossing the original *cca1-1* in Wassilewskija (Ws) background (Green and Tobin, 1999) six times into Col-0 (Yakir *et al*., 2009). Primers for characterizing *cca1-1* were described previously (Green and Tobin, 1999). Primers for characterizing *ataf2-2* were designed using the web tool provided by the Salk Institute (http://signal.salk.edu/tdnaprimers.2.html). The p*BAS1*:BAS1-GUS and p*SOB7*:SOB7-GUS constructs and the histochemical GUS staining procedures were described previously (Sandhu *et al*., 2012; Peng *et al*., 2015). Plant GUS-staining images were photographed using the Leica MZ10 F modular stereo microscope and the Leica DFC295 digital microscope color camera. For transgenic events, homozygous single-locus T-DNA insertion lines were selected for cross and further analysis. Unless otherwise stated, all seeds were surface-sterilized by ethanol, plated on half-strength Linsmaier and Skoog medium with 10 g/L phytagel (Sigma-Aldrich, made in USA) and 15 g/L sucrose, put for stratification at 4 °C in the dark for four days, and grown in growth chambers at 25 °C in 80 μmol m^−2^ s^−1^ continuous white light (red:far-red light ratio 1:1). Unless otherwise stated, four-day-old seedlings were used for total RNA extraction and hypocotyl measurements. For circadian analysis of gene expression, seedlings were grown in a 12-h light and 12-h dark photoperiod for seven days before RNA samples were extracted at 4-h intervals. Seedlings continued to grow in the same photoperiod during the two-day RNA extraction schedule. For seed collection, seedlings were transferred to the greenhouse and grown at 22 °C in 16 hours of light and 8 hours of dark photoperiod. Seeds for all physiological and molecular assays are from plants grown at the same time under the same conditions.

### DNA-protein and protein-protein interaction assays

The Gateway-compatible yeast one-hybrid (Y1H) system used in this research was developed by Deplancke *et al*. (2006). the promoter DNA fragments (baits) were amplified using primer pairs with adaptor sequences of the attB4 and attB1R sites, respectively (Deplancke *et al*., 2004). The baits were cloned into pDONR-P4-P1R (Invitrogen) via BP reactions (Gateway BP Clonase II, Invitrogen). The resulting pDONR-bait constructs were used for LR reactions with Y1H destination vector pMW#2 (Gateway LR Clonase II, Invitrogen). pMW#2 contains the Gateway cassette of attR4 and attL1 recombination sites and a *HIS3* (pMW#2) reporter gene. The resulting pMW#2-bait constructs were digested by *Xho*I to be linearized. Then DNA bait::*HIS3* (pMW#2-bait) sequences were integrated via homologous recombination into the mutant *HIS3* locus of the yeast strain YM4271 developed for Y1H analysis. The successful integrations of baits in yeast genomes were verified by PCR using the combinations of bait- and vector-specific primers (Deplancke *et al*., 2006). The self-activation of HIS3 was tested by yeast tolerance to gradient concentrations (0 - 80 mM) of 3-AT (3-aminotriazole, a competitive inhibitor of the His3p enzyme). After self-activation tests of HIS3 reporters, the yeast bait clones with the lowest background of reporter activity (self-activation) were selected and used to test their interactions with the preys. The sequences of baits p*BAS1* -EE, p*SOB7*-CBS, p*BAS1*-CBS1, p*BAS1*-EEm and p*SOB7*-CBSm were described previously (Peng *et al*., 2015). The sequence of bait p*ATAF2*-CBS is listed in Table S1. The full-length cDNA clone of *CCA1* (C105127; AT2G46830.2) was obtained from ABRC. *CCA1* was cloned into the Gateway-compatible prey vector pACT2-GW (pACT2-GW-CCA1) and its interaction with the baits mentioned above was tested. An empty prey vector was used as a negative control. The procedure of the yeast two-hybrid (Y2H) assay was described previously (Zhao *et al*., 2013). *ATAF2* cDNA was cloned into the Gateway-compatible bait vector pBTM116-D9 (pBTM116-D9-ATAF2). The prey construct pACT2-GW-CCA1 was used to transform yeast strain A. After testing for self-activation, the resulting clone was used for transformation of the bait construct pBTM116-D9-ATAF2. The empty bait vector was used as a negative control. The CCA1-ATAF2 interaction was tested by yeast tolerance to 3- AT and ability to grow in SDIV media deprived of uracil, histidine, leucine and tryptophan. The PCR-amplified sequences in all constructs used in this research were verified by sequencing.

### Transcript analysis

Transcript accumulations of *BAS1*, *SOB7*, *ATAF2* and *CCA1* were measured by qRT-PCR. Total RNA was extracted from four-day-old seedlings using the RNeasy Plant Kit (Qiagen). The DNase I Digestion Set (Sigma) was used to perform on-column elimination of genomic DNA contamination. First-strand cDNA was synthesized using the iScript Reverse Transcription Supermix for RT-qPCR (Bio-Rad). Quantitative PCRs (qPCRs) were performed using the SsoAdvanced Universal SYBR Green Supermix (Bio-Rad) and the CFX96 Real-Time System (Bio-Rad). The Bio-Rad CFX Manager software was used to analyze and compare data using the ΔΔC_T_ method. Relative expression levels of target genes were determined by normalizing to the transcript levels of *UBQ10*. Each data point represents nine replicates (three biological replicates × three technical replicates). qPCR primers for *UBQ10*, *ATAF2*, *BAS1* and *SOB7* were described previously (Peng *et al*., 2015). qPCR primers for *CCA1* are 5’-TCGAAAGACGGGAAGTGGAACG-3’ and 5’-GTCGATCTTCATTGGCCATCTCAG-3’. All qPCR primers were designed using QuantPrime (http://quantprime.mpimp-golm.mpg.de/; Arvidsson *et al*., 2008).

### Hypocotyl measurements

Seed plating and hypocotyl measurement were described previously (Favero *et al*., 2016; 2017). For brassinolide (BL) treatment assays, seeds were put on BL-containing plates from the beginning of the experiments. The same volume of ethanol was used to dissolve gradient concentrations of BL and added to the media including the non-BL control. All four-day-old seedlings were scanned/photographed and measured using NIH ImageJ (Schneider *et al*., 2012a). Each data point represents the result of thirty tallest seedlings. All experiments were repeated three times. Each independent experiment showed the similar trend of differences.

### Accession numbers

Arabidopsis Genome Initiative numbers for the genes used in this study are as follows: CCA1 (AT2G46830), ATAF2 (AT5G08790), BAS1 (AT2G26710), SOB7 (AT1G17060), UBQ10 (AT4G05320).

## Results

### CCA1 binds the EE and CBS motifs on BAS1 and SOB7 promoters, respectively

ATAF2 binds three EE- and CBS-containing fragments of the *BAS1* and *SOB7* promoters (Fig. 1A), including p*BAS1*-EE (−731 to −504), p*BAS1*-CBS1 (−844 to −786) and p*SOB7*-CBS (−1623 to −1524) (Peng *et al*., 2015). Since both EE and CBS elements are known binding targets of CCA1 (Pan *et al*., 2009), we tested, via targeted Y1H assays, the binding capability of CCA1 to the three *BAS1* and *SOB7* promoter fragments mentioned above. CCA1 was confirmed to interact with p*BAS1*-EE (Fig. 1B) and p*SOB7*-CBS (Fig. 1C). Unlike ATAF2, CCA1 did not bind p*BAS1*-CBS1 in our assay (Fig. 1D).

**Fig. 1.**
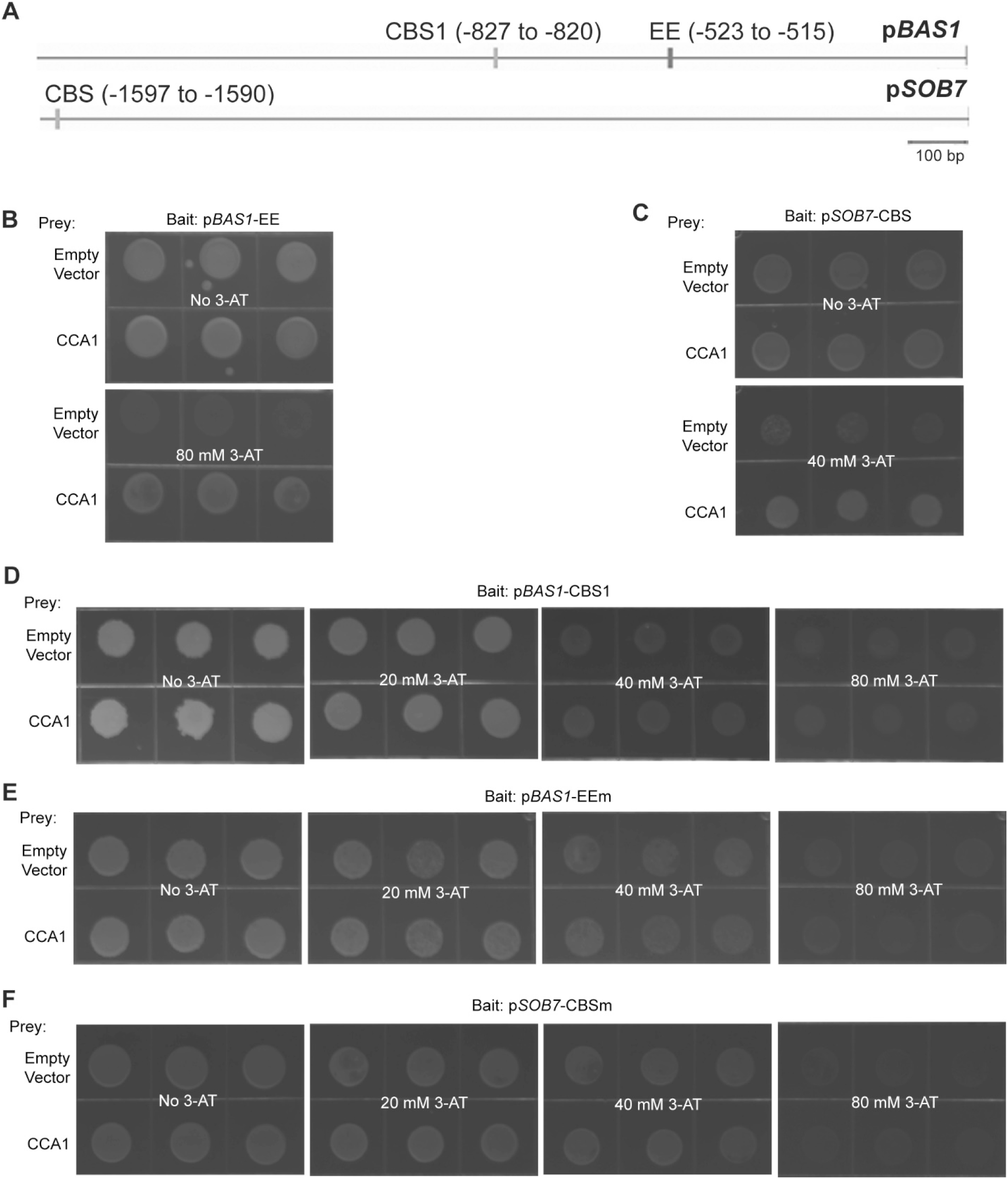
CCA1 binds the EE and CBS motifs on *BAS1* and *SOB7* promoters, respectively. (A) The *BAS1* promoter harbors both EE and CBS motifs, while only one CBS motif exists in the *SOB7* promoter. CCA1 interacted with p*BAS1*-EE (B) and p*SOB7*-CBS (C), but not with p*BAS1*-CBS1 (D) in targeted Y1H assays. CCA1 did not interact with p*BAS1*-EEm (E) or p*SOB7*-CBS (F), in which the EE or CBS motifs have been mutated, respectively. For each Y1H interaction tested, the indicated bait sequence was integrated into the mutant *HIS3* locus of the yeast strain YM4271. The bait-integrated yeast clone with lowest self-activation was transformed with the indicated prey construct and empty prey vector (negative control), and then plated on selection medium supplemented with 3-AT of described concentrations. Yeast clones were grown at 28 °C for three to four days. Three independent clones were shown for each sample.

Two EE/CBS-mutated fragments (Peng *et al*., 2015), p*BAS1*-EEm (EE was mutated from AAAATATCT to AACATATCT) and p*SOB7*-CBSm (CBS was mutated from AGATTTTT to AGATTCTT), were used to test whether the interactions between CCA1 and p*BAS1*-EE/p*SOB7*-CBS were mediated by the EE and CBS motifs, respectively. Both p*BAS1*- EEm (Fig. 1E) and p*SOB7*-CBSm (Fig. 1F) lost their binding capacity to CCA1 in targeted Y1H assays, indicating that the EE and CBS motifs are responsible for the binding of CCA1 to *BAS1* and *SOB7* promoters, respectively.

### CCA1 is a repressor of BAS1-GUS and SOB7-GUS activity

To test the effects of CCA1 on BAS1 and SOB7 activity, two constructs p*BAS1*:BAS1-GUS and p*SOB7*:SOB7-GUS (genomic DNA translational fusions with 1.6 and 2.1 kb of their native promoters, respectively; Sandhu *et al.*, 2012; Peng *et al.*, 2015) were used to transform the *CCA1* loss-of-function mutant *cca1-1*. Approximately 25% of the T1 primary transformants of both p*BAS1*:BAS1-GUS/*cca1-1* (Fig. 2A) and p*SOB7*:SOB7-GUS/*cca1-1* (Fig. 2B) conferred a severe dwarf phenotype associated with BR deficiency (BR-dwarf). Similar BR-dwarf transformants were observed when expressing the two constructs in the *ATAF2* loss-of-function mutant *ataf2-2*, while none of the p*BAS1*:BAS1-GUS and p*SOB7*:SOB7-GUS transgenic plants in Col-0 background showed dwarfism (Peng *et al*., 2015). The results indicate that like ATAF2, CCA1 may also suppress the expression and activity of *BAS1* and *SOB7*.

**Fig. 2.**
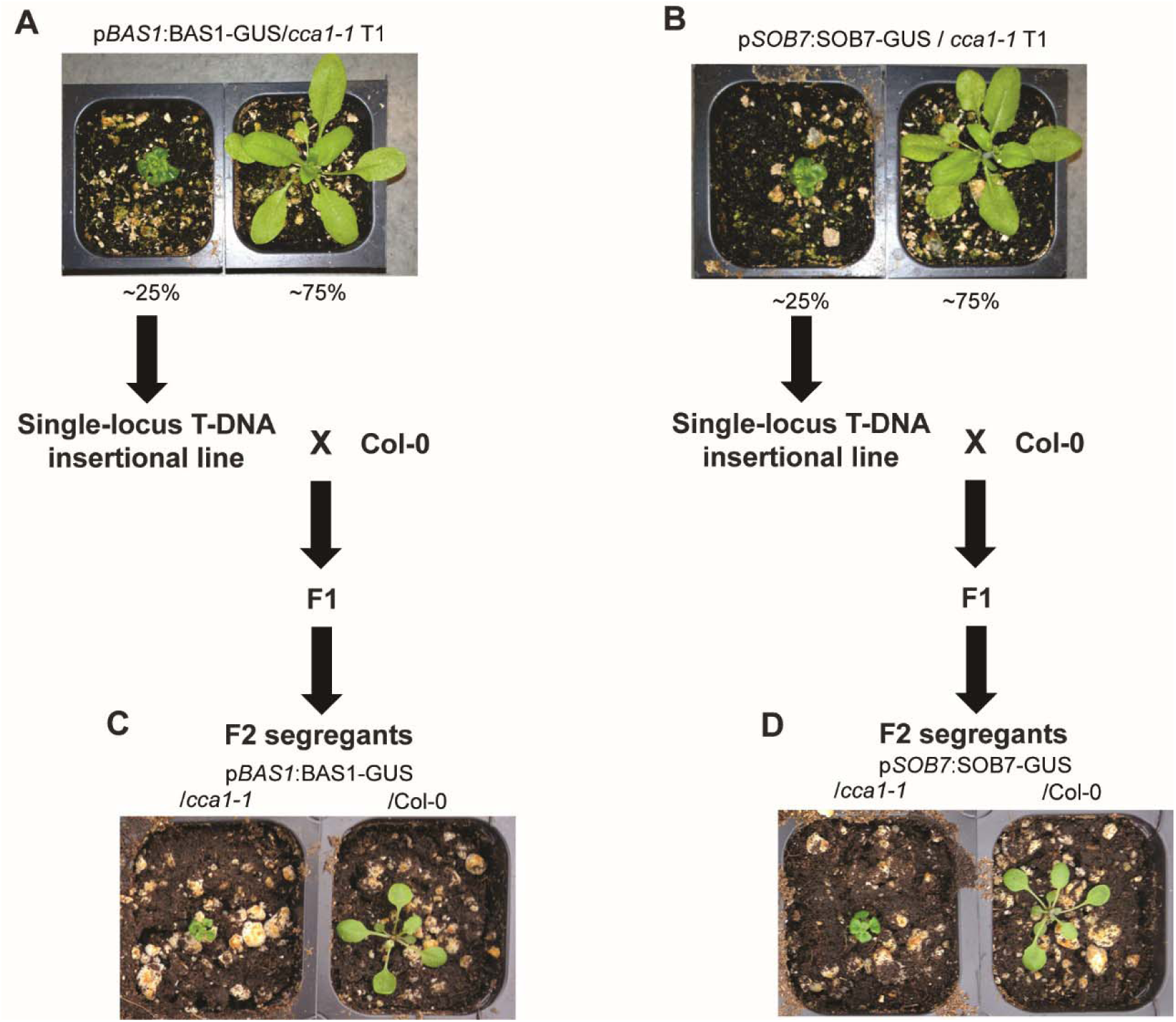
CCA1 is a repressor of *BAS1* and *SOB7* expression. Ectopic expression of p*BAS1*:BAS1- GUS (A) and p*SOB7*:SOB7-GUS (B) in the *cca1-1* background caused BR deficiency-associated dwarfism in about one-fourth T1 plants of both transgenic events. Single-locus T-DNA insertional p*BAS*1:BAS1-GUS/*cca1-1* and p*SOB7*:SOB7-GUS/*cca1-1* homozygous T3 lines were selected from BR-dwarf plants and crossed with Col-0, respectively. Homozygous p*BAS*1:BAS1-GUS (C) and p*SOB7*: SOB7-GUS (D) sibling lines in *cca1-1* and Col-0 backgrounds were selected from the two F2 segregation populations for comparison of morphology.

To compare the activity of p*BAS1*:BAS1-GUS or p*SOB7*:SOB7-GUS in wild-type (Col-0) and *cca1-1* backgrounds with identical insertion sites in the *Arabidopsis* genome, we adopted a cross-segregation approach previously applied to *ataf2-2* (Peng *et al*., 2015). Homozygous T3 plants were isolated from the BR-dwarf p*BAS1*:BAS1-GUS/*cca1-1* and p*SOB7*:SOB7-GUS/*cca1-1* lines with T-DNA inserted at a single locus. Those homozygous T-DNA insertional plants were crossed with Col-0, and the F2 segregants were genotyped. Many p*BAS1*:BAS1-GUS/*cca1-1* and p*SOB7*:SOB7-GUS/*cca1-1* F2 segregants retained the BR-dwarf phenotype, whereas all p*BAS1*:BAS1-GUS/Col-0 and p*SOB7*:SOB7-GUS/Col-0 siblings were morphologically normal (Fig. 2C, D). The results confirmed that the BR-dwarf phenotype of p*BAS1*:BAS1-GUS/*cca1-1* and p*SOB7*:SOB7-GUS/*cca 1-1* transgenic plants were caused by the disruption of CCA1.

### CCA1 modulates the tissue-specific protein accumulation patterns of BAS1-GUS and SOB7-GUS

Both BAS1-GUS and SOB7-GUS have specific accumulation patterns that limit their presence in certain tissues of seedlings and plant organs (Sandhu *et al*., 2012). BAS1-GUS accumulates in seedling roots, the shoot apex, and certain leaf regions, whereas SOB7-GUS activity can only be observed in the root tip and elongation zone (Peng *et al*., 2015). Using CCA1-GUS transgenic lines, Pruneda-Paz *et al*. (2009) revealed that CCA1 exhibits expression throughout the whole seedling except the roots. Based on our previous results (Figs. 1 and 2), CCA1 may act as a tissue-specific repressor of BAS1 and SOB7. To test this hypothesis, we performed GUS staining on F3 homozygous segregants of p*BAS1*:BAS1-GUS/Col-0 (Fig. 3A-E), p*BAS1*:BAS1-GUS/*cca1-1* (Fig. 3F-J), p*SOB7*:SOB7-GUS/Col-0 (Fig. 3K-O), and p*SOB7*:SOB7-GUS/*cca1-1* (Fig. 3P-T). Five-day-old seedlings, cauline and rosette leaves, as well as flowers and siliques were stained (Fig. 3). The results showed that BAS1 and SOB7 expression expanded to more tissues with the disruption of *CCA1*. In a *cca1-1* background, both BAS1 and SOB7 GUS-fusion signals were dramatically expanded and enhanced in seedlings, leaves, flowers and siliques when compared with their expression patterns in the wild type (Col-0) (Fig. 3). BAS1 and SOB7 expression was also found in stigma and peduncle tissues when *CCA1* was disrupted (Fig. 3D, I, N, S). The disruption of *CCA1* did not significantly alter BAS1 or SOB7 expression patterns in seedling roots (Fig. 3A, F, K, P). This result is consistent with the observation that CCA1 had no visible expression in the root tissues of light-grown seedlings (Pruneda-Paz *et al*., 2009).

**Fig. 3.**
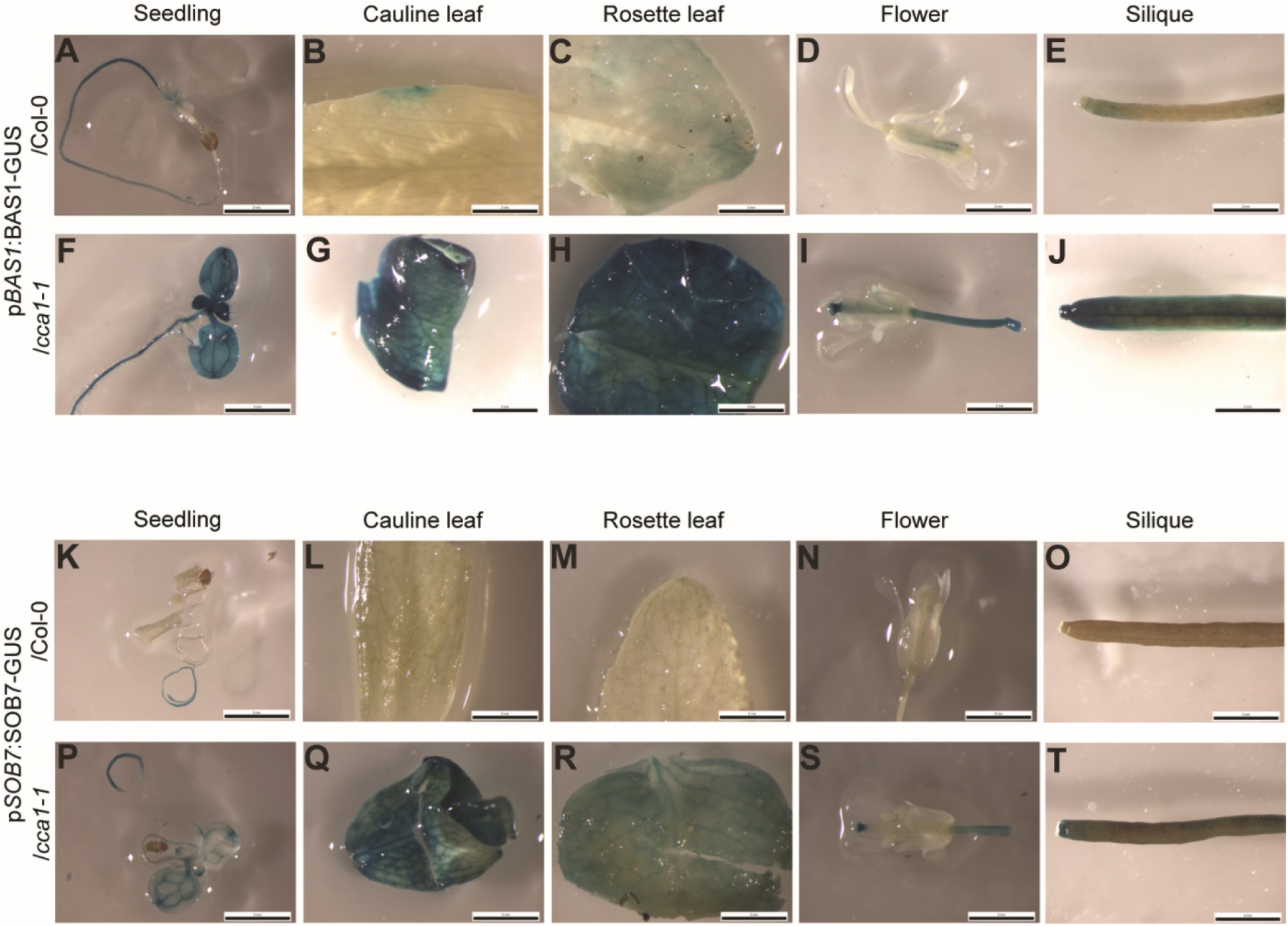
CCA1 modulates the tissue-specific expression patterns of BAS1 and SOB7. The role of CCA1 in restricting BAS1 and SOB7 expression within certain tissues was demonstrated by GUS analysis on F3 homozygous segregants of p*BAS1*:BAS1-GUS/Col-0 (A-E), p*BAS1*:BAS1-GUS/*cca1-1* (F-J), p*SOB7*:SOB7-GUS/Col-0 (K-O), and p*SOB7*:SOB7-GUS/*cca1-1* (P-T). Scale bars: 2 cm.

### CCA1 and ATAF2 differentially suppress the transcript accumulation of BAS1 and SOB7

Since CCA1 and ATAF2 have similar functions in suppressing BAS1-GUS and SOB7-GUS accumulation, we made the *cca1-1 ataf2-2* double mutant and compared it to the single mutants and the wild type with regard to *BAS1* and *SOB7* transcript accumulation. For four-day-old seedlings grown at 25 °C in 80 μmol m^−2^ s^−1^ continuous white light or darkness, *cca1-1*, *ataf2-2* and the *cca1-1 ataf2-2* double mutant showed similarly elevated *BAS1* expression when compared to the wild type (Col-0) (Fig. 4A, B), demonstrating that the removal of either *CCA1* or *ATAF2* is sufficient to de-repress *BAS1* transcript accumulation. In contrast, in white light the *cca1-1 ataf2-2* double mutant conferred significantly higher *SOB7* transcript accumulation than either *cca1-1* or *ataf2-2* single mutants (Fig. 4C). However, in darkness the genetic impact of *CCA1* or *ATAF2* on *SOB7* transcript accumulation is similar to that of *BAS1* (Fig. 4D). These results indicate that CCA1 and ATAF2 can additively suppress *SOB7* transcript accumulation in white light but not in darkness.

**Fig. 4.**
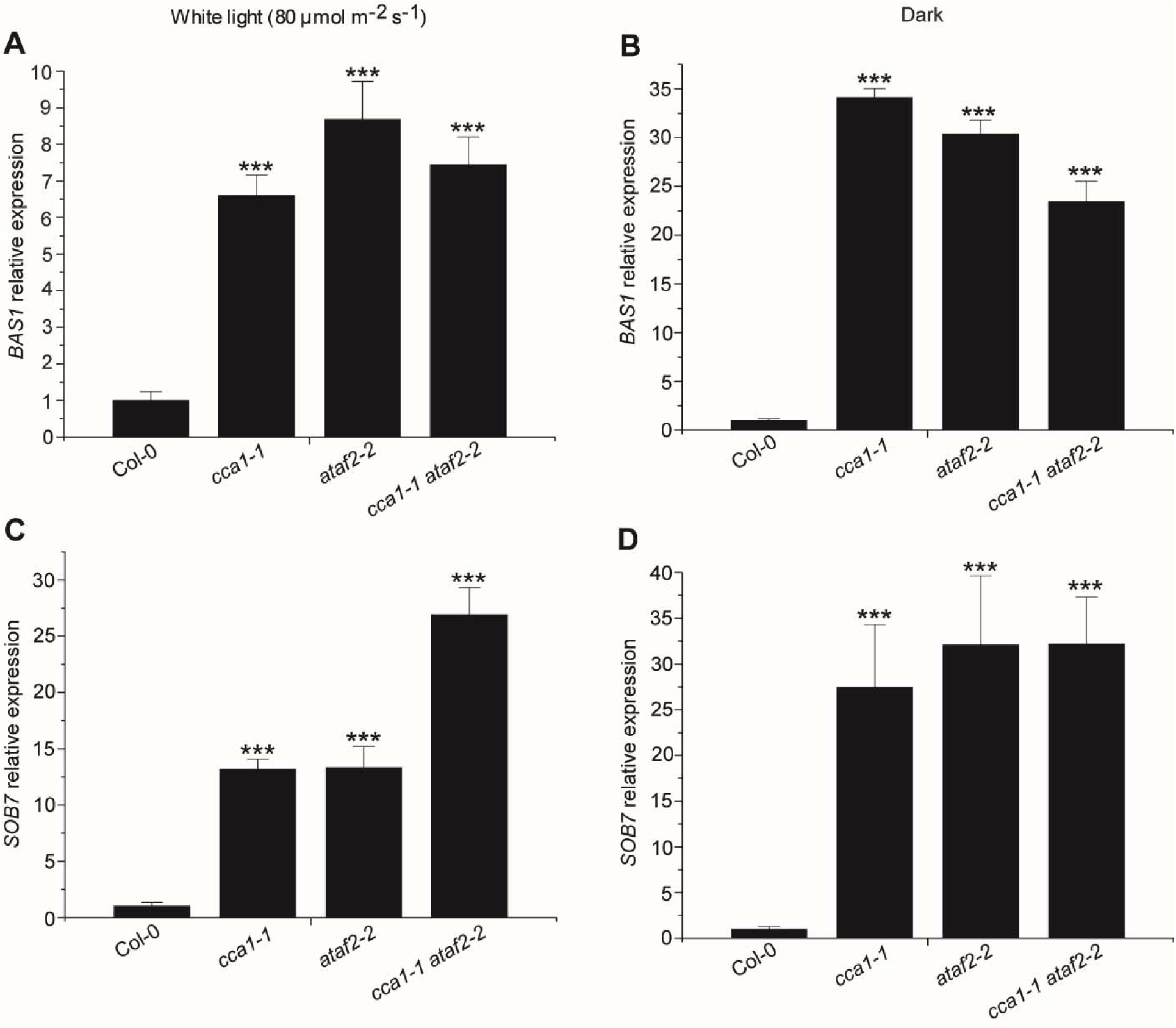
CCA1 and ATAF2 differentially suppress the transcript accumulation of *BAS1* and *SOB7*. For four-day-old seedlings grown at 25 °C in 80 μmol m^−2^ s^−1^ continuous white light (A) or darkness (B), *cca1-1*, *ataf2-2* and the *cca1-1 ataf2-2* double mutant showed similarly elevated *BAS1* expression when compared to the wild type (Col-0). In contrast, in white light the *cca1-1 ataf2-2* double mutant conferred significantly higher *SOB7* transcript accumulation than either *cca1-1* or *ataf2-2* single mutants (C). However, in darkness the genetic impact of *CCA1* or *ATAF2* on *SOB7* transcript accumulation is similar to that of *BAS1* (D). Each qRT-PCR value is the mean of results from three biological replicates × three technical replicates (*n*=9). Error bars denote the s.e.m. \*\*\**P*<0.001.

### CCA1 and ATAF2 both impact seedling responsiveness to exogenous BL

Since BRs promote hypocotyl growth in white light but have the opposite, suppressing, effect under darkness (Turk *et al*., 2003; Peng *et al*., 2015), *Arabidopsis* seedlings with elevated *BAS1* or *SOB7* expression are less responsive to exogenous BRs when compared to the wild type (Turk *et al*., 2005; Peng *et al*., 2015). To test the BR sensitivity of Col-0, *cca1-1*, *ataf2-2*, and *cca1-1 ataf2-2*, seedlings of all four genotypes were grown on media with gradient concentrations of BL (0, 10, 100 and 1000 nM) for four days in 80 μmol m^−2^ s^−1^ white light and darkness, respectively. Since *cca1-1* had slightly shorter hypocotyls than Col-0 even without BL treatment (Fig. 5A), seedling hypocotyl lengths in response to exogenous BL were described as percent changes instead of their absolute values (Fig. 5B). The hypocotyl growth of both *cca1-1* and *ataf2-2* was less responsive to BL treatments when compared with that of Col-0 (Fig. 5B). *cca1-1 ataf2-2* was even more insensitive to BL than the two single mutants (Fig. 5B). Based on our previous gene expression results (Fig. 4), the additive effect of CCA1 and ATAF2 in regulating BR-responsive hypocotyl growth in light is due to their collaborative suppression of *SOB7* expression. In contrast, *cca1-1 ataf2-2* did not show higher BL insensitivity than either *cca1-1* or *ataf2-2* in the dark (Fig. S1).

**Fig. 5.**
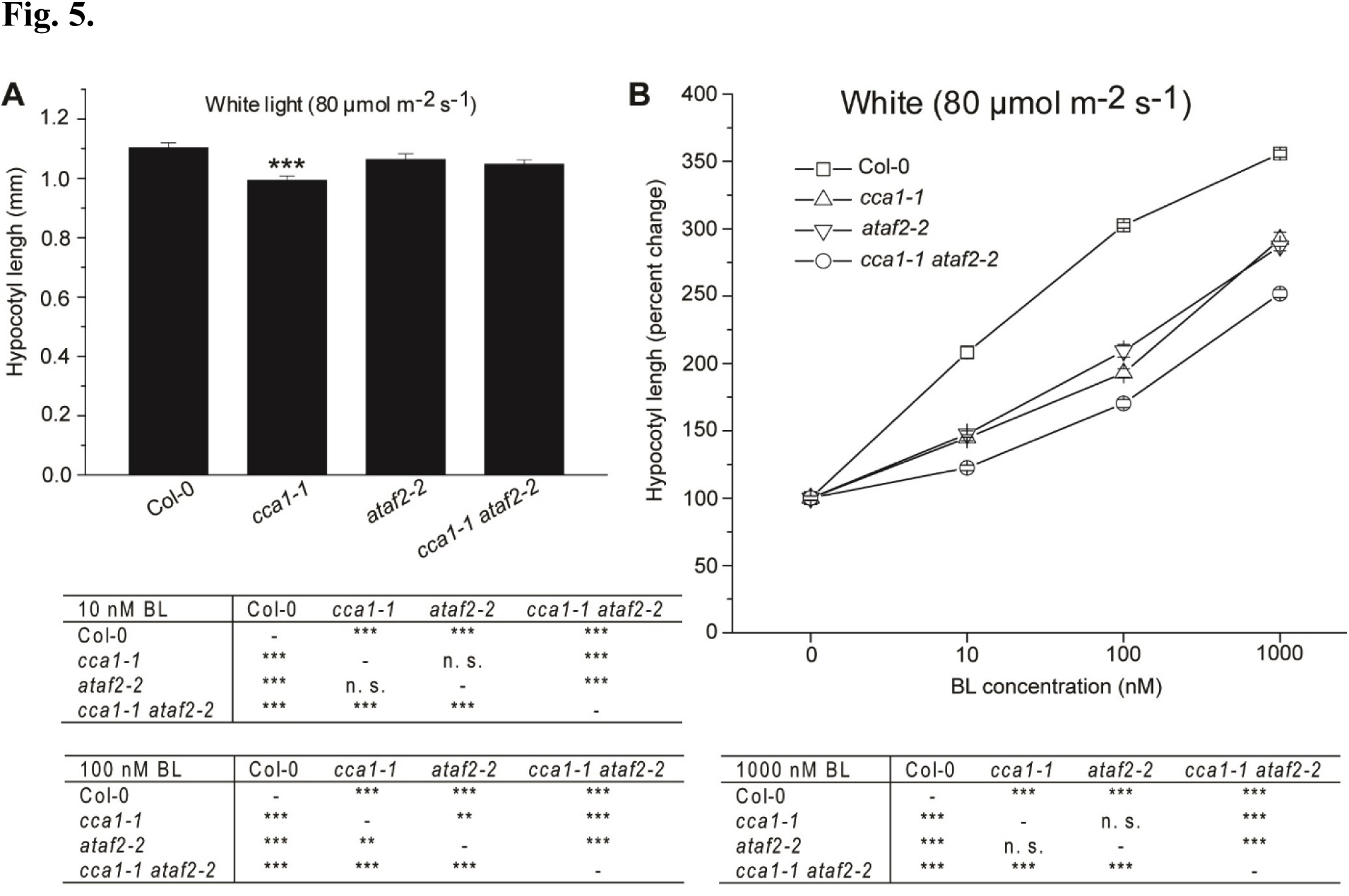
*cca −1 ataf2-2* is less responsive to exogenous BL treatments than either single mutant under light. (A) *cca −1* had slightly shorter hypocotyls than Col-0. (B) The hypocotyl growth of both *cca1-1* and *ataf2-2* was less responsive to BL treatments when compared with that of Col-0. *cca1-1 ataf2-2* was even more insensitive to BL than the two single mutants. Four-day-old seedlings grown at 25 °C in 80 μmol m^−2^ s^−1^ continuous white light were used for hypocotyl measurements. Each data point represents the average of measurements from 30 seedlings (*n*=30). Error bars denote the standard deviations. n. s. *P*>0.01, \*\**P*<0.01, \*\*\**P*<0.001.

### CCA1 is not feedback regulated by BRs

In addition to being a repressor of the BR-inactivating genes *BAS1* and *SOB7*, *ATAF2* is transcriptionally suppressed by exogenous BL, which forms a feedback regulatory loop (Peng *et al*., 2015). This led to the hypothesis that *CCA1* may also be feedback regulated by exogenous BL. When treated with BL, *CCA1* transcript accumulation in Col-0 did not show any significant change (Fig. 6). The result indicated that unlike *ATAF2*, *CCA1* is not subject to BR-mediated transcriptional feedback regulation.

**Fig. 6.**
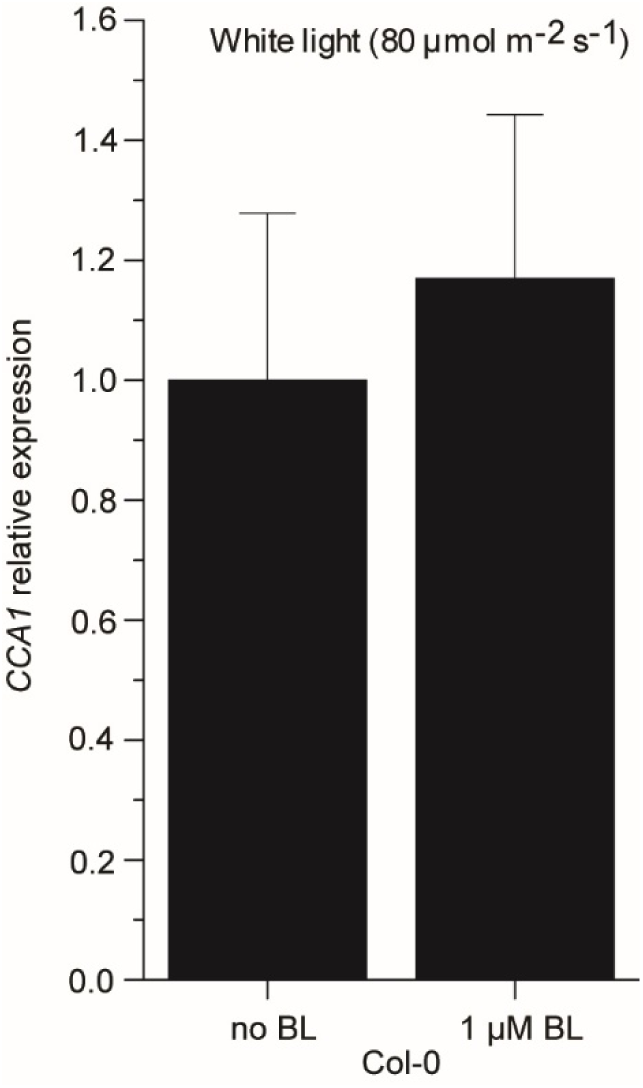
*CCA1* is not feedback regulated by BRs. *CCA1* transcript accumulations in Col-0 did not show any significant change when treated with 1 μM BL. Four-day-old seedlings grown at 25 °C in 80 μmol m^−2^ s^−1^ continuous white light were used for RNA extraction. Each qRT-PCR value is the mean of results from three biological replicates × three technical replicates (*n*=9). Error bars denote the s.e.m.

### CCA1 interacts with ATAF2 at both the DNA-protein and protein-protein levels

Since CCA1 and ATAF2 share the same binding sites (EE and CBS) (Peng *et al*., 2015; Fig. 1), and both act as repressors of the BR-inactivating genes *BAS1* and *SOB7* (Peng *et al*., 2015; Figs. 2-4), we tested whether these two TF genetically or physically interact. Like *BAS1* and *SOB7*, the *ATAF2* promoter also contains a CBS motif (−577 to −570; Fig. 7A), which is a potential binding target for CCA1. A 63-bp *ATAF2* promoter fragment, p*ATAF2*-CBS (−598 to −536; Table S1), was used as the bait in a targeted Y1H assay to test its interaction with CCA1. CBS is the only predicted TF binding site harbored by p*ATAF2*-CBS. The Y1H result demonstrates CCA1 can bind to the promoter of *ATAF2* (Fig. 7B). Compared to Col-0, *cca1-1* seedlings showed significantly higher *ATAF2* transcript accumulation when grown in continuous white light (Fig. 7C), whereas the opposite trend was observed in dark-grown seedlings (Fig. 7D). The results above reveal that CCA1 can act as either a repressor or an activator of *ATAF2* expression depending on the light conditions. On the other hand, no significant changes of *CCA1* expression in *ataf2-2* seedlings were observed in either continuous light (Fig. S2A) or dark condition (Fig. S2B), which indicates ATAF2 is not a transcriptional regulator of *CCA1*. Furthermore, CCA1 physically interacts with ATAF2 in a targeted Y2H assay (Fig. 7E).

**Fig. 7.**
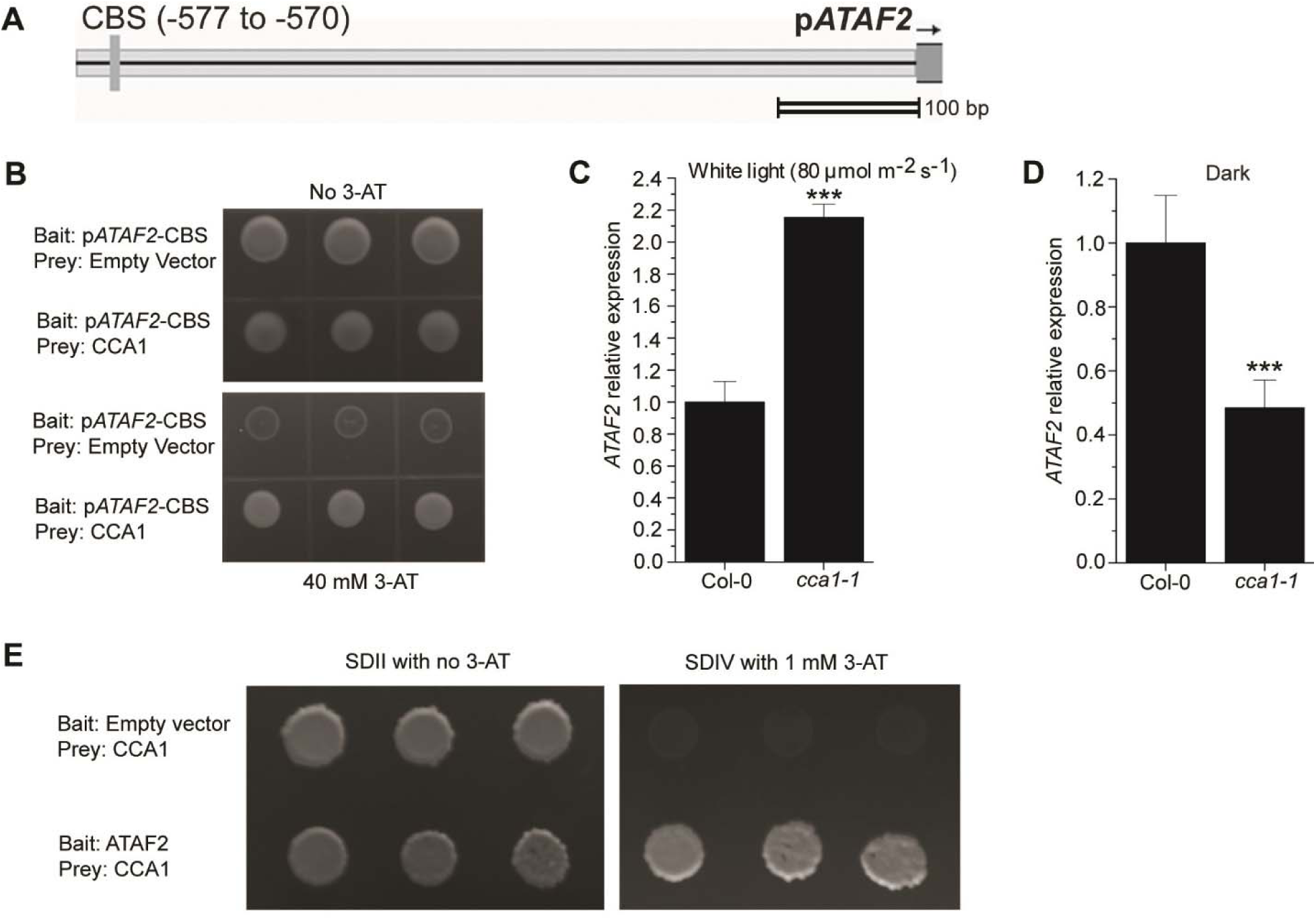
CCA1 interacts with ATAF2 at both DNA-protein and protein-protein levels. (A) The *ATAF2* promoter contains a CBS motif. (B) CCA1 binds p*ATAF2*-CBS in a targeted Y1H assay. Compared to Col-0, *cca1-1* seedlings showed significantly higher *ATAF2* transcript accumulation when grown in continuous white light (C), whereas opposite trend was observed in dark-grown seedlings (D). (E) CCA1 physically interacts with ATAF2 in a targeted Y2H assay. For Y1H assay, the p*ATAF2*-CBS bait was integrated into the mutant *HIS3* locus of the yeast strain YM4271. The bait-integrated yeast clone with lowest self-activation was transformed with the CCA1 prey construct and empty prey vector (negative control). The interaction between CCA1 and p*ATAF2*-CBS was tested by yeast tolerance to 3-AT. For Y2H assay, the CCA1 prey construct was used to transform yeast strain A. After testing for self-activation, the resulting clone was used for transformation of the ATAF2 bait construct and the empty bait vector (negative control). The CCA1-ATAF2 interaction was tested by yeast tolerance to 3-AT and the ability to grown on SDIV media. All yeast clones were grown at 28 °C for three to four days. Three independent clones were shown for each Y1H or Y2H sample. Four-day-old seedlings grown at 25 °C in 80 μmol m^−2^ s^−1^ continuous white light were used for RNA extraction. Each qRT-PCR value is the mean of results from three biological replicates × three technical replicates (*n*=9). Error bars denote the s.e.m. \*\*\**P*<0.001.

### ATAF2, BAS1 and SOB7 are all subject to circadian regulation

Since CCA1 acts as a core regulator for circadian clock (Wang and Tobin, 1998), we tested whether *ATAF2*, *BAS1* and *SOB7* expression are circadian regulated in wild-type (Col-0) Arabidopsis seedlings (Fig. 8). After seedlings were grown in 12-h/12-h of light/dark cycle for seven days, gene expression was monitored at 4-h intervals for two days under the same photoperiod setting. The results indicated that *ATAF2*, *BAS1* and *SOB7* are all circadian-regulated with distinct expression patterns (Fig. 8). Transcript accumulation of *ATAF2* kept decreasing in the dark period and began to increase after transiting to light (Fig. 8). In contrast, both *BAS1* and *SOB7* showed higher expression levels in the dark than under light, and their transcript accumulation peaks appeared after entering the dark period for four hours (Fig. 8).

**Fig. 8.**
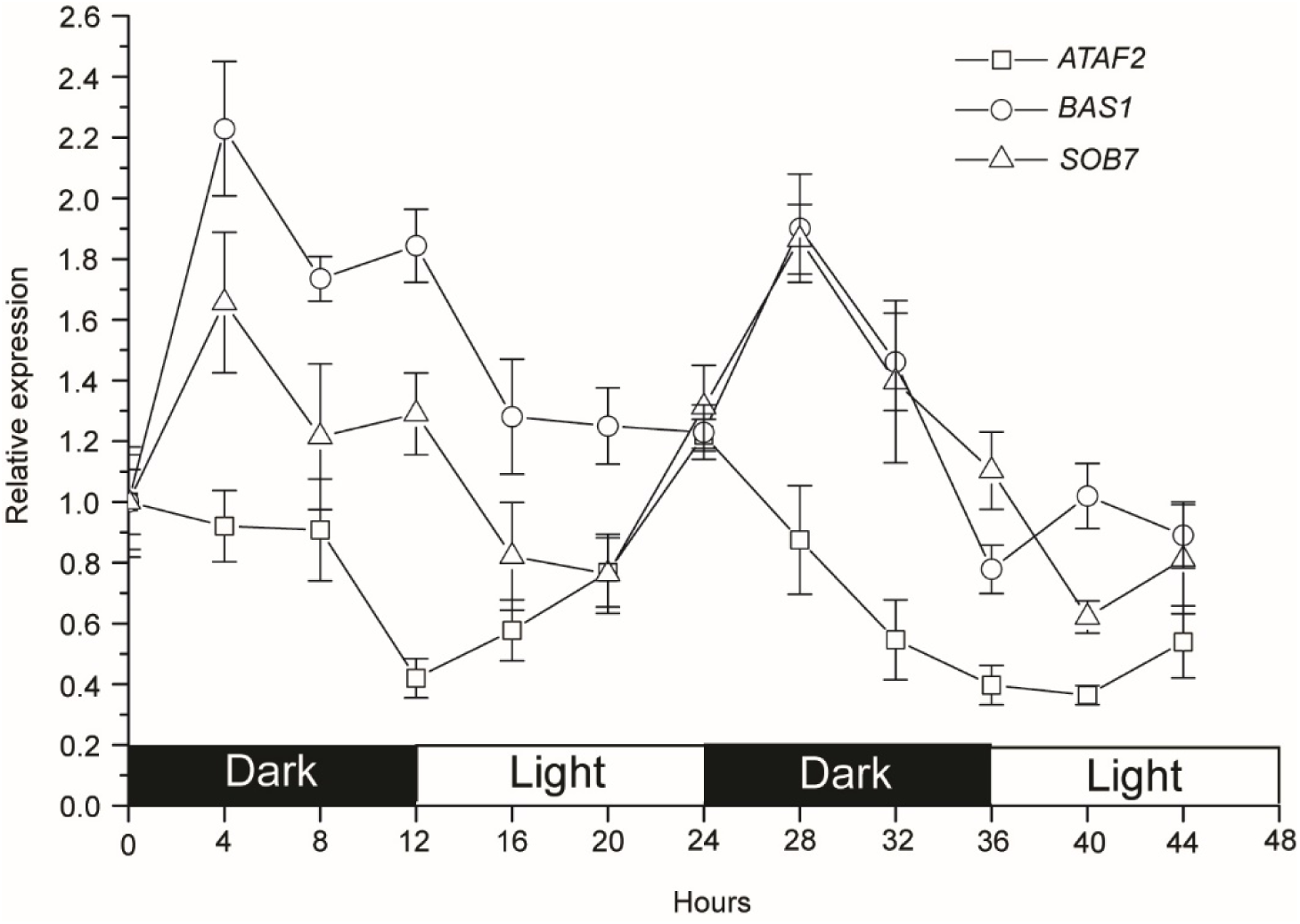
*ATAF2*, *BAS1* and *SOB7* are all circadian-regulated with distinct expression patterns. Transcript accumulation of *ATAF2* kept decreasing in the dark period and began to increase after transiting to light. In contrast, both *BAS1* and *SOB7* showed higher expression levels in the dark than under light, and their transcript accumulation peaks appeared after entering the dark period for four hours. After Col-0 seedlings were grown in 12-h/12-h of light/dark cycle for seven days, gene expression was monitored at 4-h intervals for two days under the same photoperiod setting. Each qRT-PCR value is the mean of results from three biological replicates × three technical replicates (*n*=9). Error bars denote the s.e.m.

## Discussion

### BAS1, SOB7 and multiple other BR-inactivating genes contribute to BR homeostasis

BR inactivation can be achieved via multiple approaches in *Arabidopsis*, including hydroxylation (Neff *et al*., 1999; Turk *et al*., 2003), glycosylation (Poppenberger *et al*., 2005; Husar *et al.*, 2011), acylation (Roh *et al*., 2012; Wang *et al*., 2012a; Schneider *et al*., 2012b; Zhu *et al*., 2013b; Choi *et al*., 2013; Zhang and Xu, 2018), and other unknown or unconfirmed mechanisms (Turk *et al*., 2005; Takahashi *et al*., 2005; Nakamura *et al*., 2005; Yuan *et al*., 2007; Marsolais *et al*., 2007; Thornton *et al*., 2010; Sandhu and Neff, 2013). At least ten BR-inactivating genes have been identified in Arabidopsis including; P450 hydroxylases, glycosyltransferases, acyltransferases, sulfotransferases and a reductase. The redundancy of BR-inactivating pathways is consistent with the fact that BRs act in local tissues at extremely low endogenous concentrations (He *et al*., 2005; Kim *et al*., 2006; Symons *et al*., 2008). The role of catabolism in maintaining BR homeostasis appears to be as critical as the biosynthesis and signaling pathways, since tissue-specific BR levels can be fine-tuned by multiple inactivating enzymes and their upstream TF regulatory cascades. For example, LOB negatively regulates BR accumulation by activating *BAS1* expression at organ boundaries (Bell *et al*., 2012). As two transcriptional repressors of *BAS1* and/or *SOB7*, ATAF2 and ARF7 integrate BR inactivation with auxin biosynthesis and signaling, seedling photomorphogenesis, disease resistance and stress tolerance (Peng *et al*., 2015; Youn *et al*., 2016).

### CCA1 is a direct repressor of both BAS1 and SOB7

The existence of EE and CBS motifs in *BAS1* and *SOB7* promoters (Peng *et al*., 2015; Fig. 1A) indicates that these two genes may be included in the regulatory network of the core circadian clock protein CCA1. The genomic approach of chromatin immunoprecipitation followed by deep sequencing (ChIP-seq), did not identify *BAS1* or *SOB7* as a target of CCA1 (Nagel *et al*., 2015; Kamioka *et al*., 2016). However, our focused analysis demonstrated that CCA1 is a direct repressor of both *BAS1* and *SOB7* (Figs. 1-4). Since P450s play critical roles in the metabolism of diverse secondary compounds, they have been used as reporters for different nodes in the circadian clock network (Pan *et al*., 2009). Therefore, it’s not surprising that both *BAS1* and *SOB7* are subject to the transcriptional regulation of CCA1. The lack of *BAS1* and *SOB7* in the genomic characterization of CCA1 targets can be explained by the inherent bias of the ChIP-seq approach in enriching highly expressed loci (Teytelman *et al*., 2013). The cause of this bias may be that DNA from actively transcribed regions tends to be more exposed to binding proteins and antibodies due to nucleosome depletion (Teytelman *et al*., 2013). Since both *BAS1* and *SOB7* have extremely low expression levels that are restricted to specific tissues (Neff *et al*., 1999; Turk *et al*., 2003; 2005; Sandhu *et al*., 2012; Peng *et al*., 2015), the two genes are more likely to be filtered out than other loci in the ChIP-seq assay.

### CCA1 regulates multiple BR signaling and metabolic genes

There have been established associations between CCA1 and BRs. Two TF-encoding genes involved in BR signaling, *AIF1* (Wang *et al*., 2009a) and *MYBL2* (Ye *et al*., 2012), have been identified by both Nagel *et al*. (2015) and Kamioka *et al*. (2016) as direct targets of CCA1. CCA1 also binds to the promoter of the BR biosynthetic gene *DWF4* and activates its expression (Zheng *et al*., 2018). This report, together with our finding that CCA1 directly suppresses the BR-inactivating genes *BAS1* and *SOB7* (Figs. 1-4), suggest that CCA1 is an overall positive regulator of BR accumulation.

### CCA1 is selective in binding EE and CBS elements

Although EE and CBS are confirmed binding motifs for CCA1, CCA1 does not associate with all of the EE or CBS in the *Arabidopsis* genome (Kamioka *et al*., 2016). This binding may require appropriate sequence context within the broader regulatory region (Kamioka *et al*., 2016). CCA1 also prefers to bind EE relative to CBS (Nagel *et al*., 2015; Kamioka *et al*., 2016). Consistent with these findings, CCA1 did not bind p*BAS1*-CBS1, but did interact with p*BAS1*- EE in our study (Fig. 1B, D, E). In contrast, ATAF2 is able to bind both p*BAS1*-EE and p*BAS1*- CBS1 (Peng *et al.*, 2015).

### CCA1 and ATAF2 have overlapping and distinct patterns in suppressing BAS1 and SOB7

Disruption of either *ATAF2* (Peng *et al*., 2015) or *CCA1* (Fig. 3A-T) led to the expansion of BAS1 and SOB7 expression to additional tissues, but the suppressing patterns of CCA1 and ATAF2 are not identical. *ATAF2* disruption triggered BAS1 and SOB7 expression in petals (Peng *et al*., 2015), whereas these changes did not exist in the *cca1-1* background (Fig. 3D, I, N, S). SOB7 expression expanded to the whole seedling leaves in *ataf2-2* (Peng *et al*., 2015), but the expansion was restricted to leaf veins in *cca1-1* (Fig. 3K, P). These tissue-specific pattern differences may reflect the distinct expression patterns of CCA1 and ATAF2.

About one-quarter of the T1 p*BAS1*:BAS1-GUS/*cca1-1* (Fig. 2A) and p*SOB7*:SOB7-GUS/*cca1-1* (Fig. 2B) transformants showed BR-dwarf phenotype. Similar BR-dwarfs were previously observed in p*BAS1*:BAS1-GUS/*ataf2-2* and p*SOB7*:SOB7-GUS/*ataf2-2* transformants (Peng *et al*., 2015). However, there is no visible dwarfism in *cca1-1*, *ataf2-2*, or the *cca1-1 ataf2-2* double mutant. Since GUS translational fusions can increases protein stability in *Arabidopsis* (Chae *et al*., 2012; Spartz *et al*., 2012), BAS1-GUS and SOB7-GUS may be more likely to confer BR-dwarfing than their native forms in the *cca1-1* and/or *ataf2-2* mutant backgrounds.

### CCA1 and ATAF2 additively suppress SOB7 expression in white light

In both light- and dark-grown seedlings, CCA1 and ATAF2 suppress *BAS1* expression without an additive effect (Fig. 4A, B). In contrast, CCA1 and ATAF2 additively suppress *SOB7* expression in seedlings grown in continuous white light (Fig. 4C). However, CCA1 and ATAF2’s suppression of *SOB7* expression is not additive in darkness (Fig. 4D). This light-dependent collaborative suppression of *SOB7* helps to explain the observation that *cca1-1 ataf2- 2* seedlings only shows greater insensitivity to exogenous BL treatments than either of the single mutants when grown in white light but not in darkness (Fig. 5; Fig. S1). Although BL is not likely to be a preferred substrate for SOB7 (Thornton, *et al.*, 2010), increased expression of SOB7 can still reduce the overall endogenous levels of BRs. It is important to note that the differential regulatory patterns on *BAS1* and *SOB7* expression are likely influenced by the binding of CCA1 and ATAF2 (Peng et al., 2015; Fig. 1), light- and tissue-specific regulation of *CCA1* and *ATAF2* expression (Wang and Tobin, 1998; Peng et al., 2015; Fig. 8), and CCA1- ATAF2 interactions at both the DNA-protein and protein-protein levels (Fig. 7). Though we have shown that CCA1 and ATAF2 physically interact via targeted Y2H analysis (Fig. 7E), attempts to test their *in planta* interaction via bimolecular fluorescence complementation did not generate positive results. Thus, CCA1-ATAF2 physical interactions *in planta* may be transient, tissue-specific or require posttranslational modification of either protein.

### Unlike ATAF2 and its family members, CCA1 is not feedback regulated by BL

As part of a feedback regulation loop, *ATAF2* expression can be suppressed by external BL treatments (Peng *et al*., 2015). Additionally, microarray data showed that three other members of the *ATAF* subfamily, *ATAF1* (*ANAC002*), *ANAC102*, and *ANAC032*, are also transcriptionally down-regulated by BL (Kleinow *et al*., 2009). In contrast, CCA1 is not feedback regulated by BL in our study (Fig. 6). Since CCA1 is a core regulator for the circadian clock, it is not surprising that BRs do not have a significant impact on its expression.

### The circadian oscillation pattern of ATAF2 is different from that of BAS1 and SOB7

*BAS1* and *SOB7* have a similar circadian oscillation pattern that shows higher expression in the dark, whereas *ATAF2*’s oscillation cycle is largely opposite with expressions decreasing in the dark and increasing in the light period (Fig. 8). This observation is consistent with our previous finding that ATAF2 is a repressor for *BAS1* and *SOB7* expression (Peng *et al.*, 2015). The expression of *CCA1* itself is also subject to circadian oscillation with peak levels occurring after dawn (Wang and Tobin, 1998). *CCA1* expression levels are higher but keep dropping in the light period and are generally lower during circadian darkness (Wang and Tobin, 1998), which is consistent with our observation that CCA1 suppresses *ATAF2* expression in the light but the effect switches to promotion in darkness (Fig. 7C, D). The comparison of oscillation patterns between *CCA1* and *BAS1/SOB7* (Wang and Tobin, 1998; Fig. 8) also support our observation that CCA1 is a repressor of *BAS1* and *SOB7* expression (Figs. 2-4).

### Light regulates ATAF2 expression via either the circadian or the photomorphogenic pathway

In seedlings grown under a 12-h light and 12-h dark photoperiod, *ATAF2* expression gradually drops in the dark and increases steadily after the transition to light, with transcript accumulation levels peaking at the beginning of the evening and being the lowest around dawn (Fig. 8). On the other hand, we previously found *ATAF2* has higher transcript accumulation in dark-grown etiolated seedlings than in seedlings grown under continuous white light and that the expression of *ATAF2* in white light is fluence-rate dependent (Peng *et al.*, 2015). *ATAF2* expression can also be suppressed when etiolated seedlings are transferred to white light (Peng *et al.*, 2015). These results indicate that *ATAF2* is transcriptionally regulated by light via either the circadian or the photomorphogenic pathway. When seedlings are grown under light/dark circadian photoperiod, *ATAF2* expression is induced during the light period. In contrast, ATAF2 has higher transcript accumulation in seedlings undergoing skotomorphogenesis than in photomorphogenic seedlings.

### Current model

We summarized the roles of CCA1 and ATAF2 in regulating BR inactivation and how circadian and photomorphogenic pathways are incorporated (Fig. 9). Both CCA1 and ATAF2 suppress the expression of the BR-inactivating genes *BAS1* and *SOB7* via direct binding to their promoters. However, the roles of CCA1 and ATAF2 with regard to BR-inactivation is dynamic with respect to the light environment. Both *BAS1* and *SOB7* are circadian regulated with higher expressions in the dark period. Transcriptionally induced by light, *CCA1* plays a role in the oscillation of *BAS1* and *SOB7*. While *ATAF2* expression is feedback suppressed by BRs, *CCA1* is not subject to this transcriptional regulation. CCA1 suppresses *ATAF2* expression in seedlings grown under light but switches to an activator for *ATAF2* in etiolated seedlings. In addition, CCA1 may also physically interact with ATAF2 at the protein level. It is important to point out, however, that this model only focuses on the components characterized in this study. Clearly, other CCA1 and ATAF2 interacting proteins, as well as additional regulatory TFs, are likely to have an impact on the overall regulation of the BR-inactivating genes *BAS1* and *SOB7*.

**Fig. 9.**
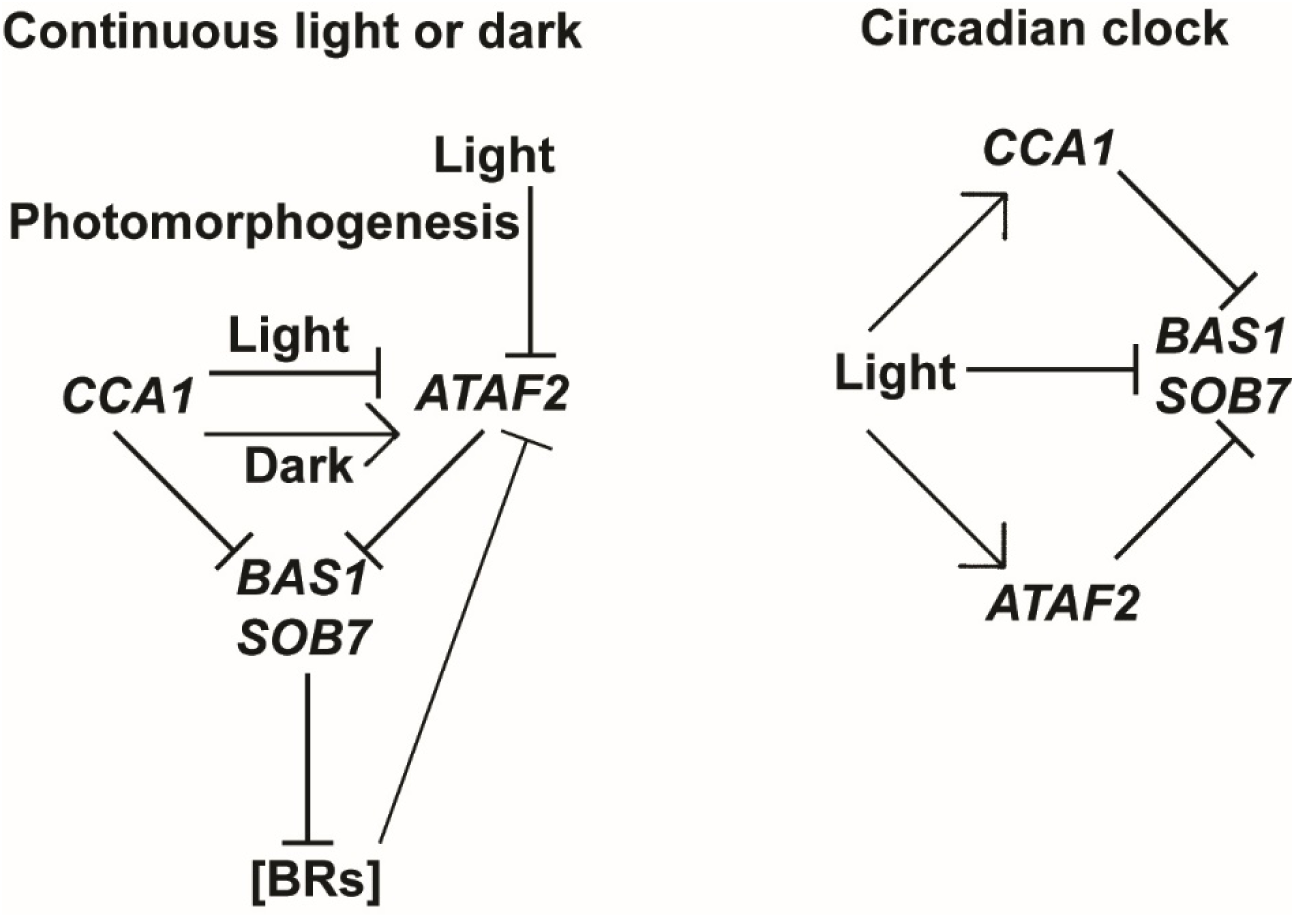
Model for the roles of CCA1 and ATAF2 in regulating BR inactivation and the incorporation of circadian and photomorphogenic pathways. Both CCA1 and ATAF2 suppress the expression of BR-inactivating genes *BAS1* and *SOB7* via direct binding to their promoters. BRs promote *Arabidopsis* hypocotyl growth under light. BAS1 and SOB7 inhibit hypocotyl elongation by catabolizing BRs. Both *BAS1* and *SOB7* are circadian regulated with higher expressions in the dark period. Transcriptionally induced by light, *CCA1* plays a role in the oscillation of *BAS1* and *SOB7*. While *ATAF2* expression is feedback suppressed by BRs, *CCA1* is not subject to such kind of transcriptional regulation. CCA1 suppresses *ATAF2* expression in seedlings grown under light but switches to an activator for *ATAF2* in etiolated seedlings. CCA1 can also physically interact with ATAF2 at protein level. Light induces *ATAF2* expression in seedlings undergoing circadian photoperiod but acts as a repressor when seedlings transit from skotomorphogenesis to photomorphogenesis.

## Acknowledgements

This research was supported by the United States National Science Foundation project 1656265 (to M.M.N.). This research was also supported by the USDA National Institute of Food and Agriculture, HATCH project 1007178 (to M.M.N.). The authors would like to thank Dr. Gaganjot Sidhu and Shahbaz Ahmad in Neff Lab for their critical review and comments on this manuscript.

## Supplemental Figure Legends

**Fig. S1.**
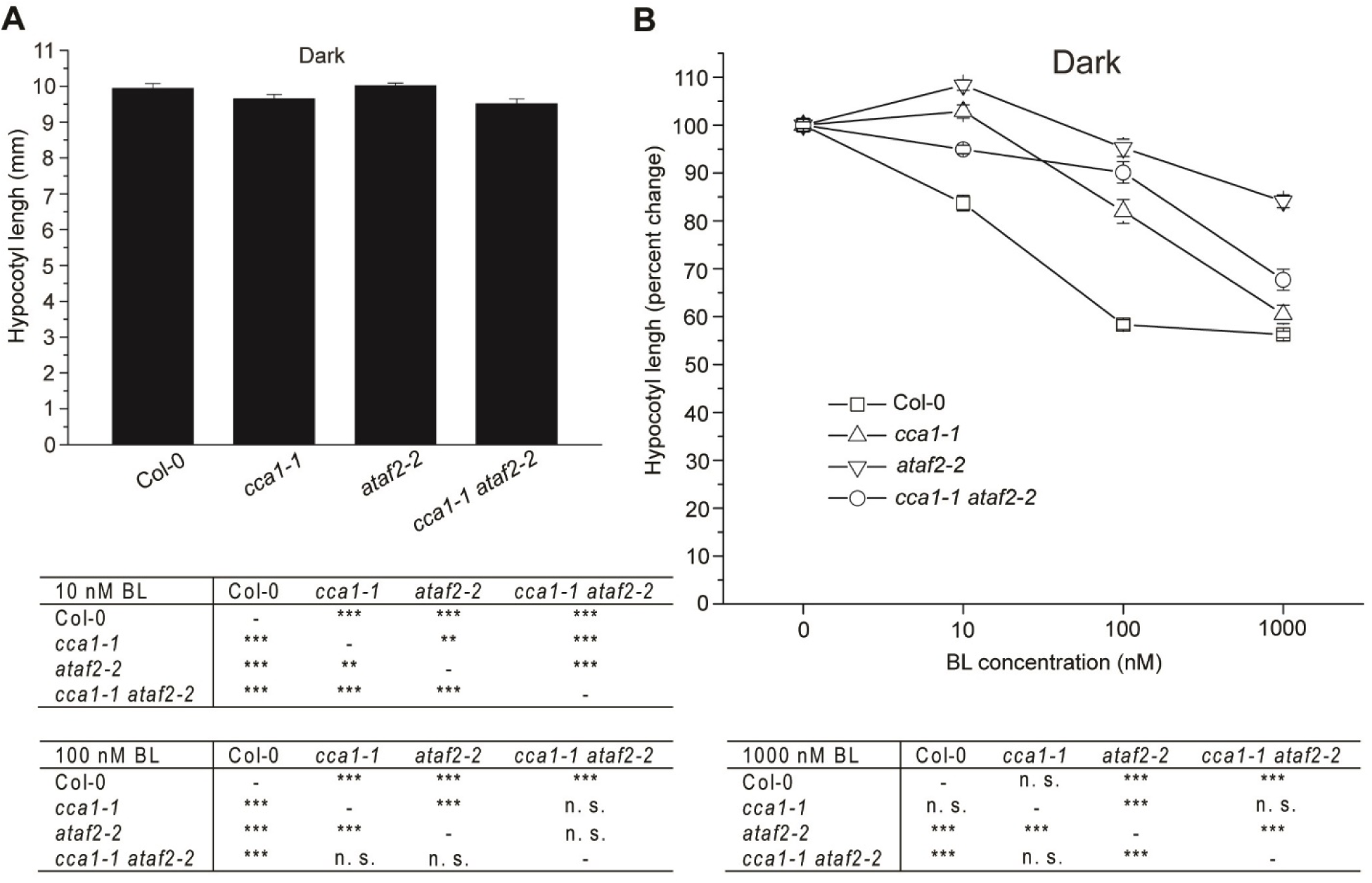
*cca1-1 ataf2-2* is not less responsive to exogenous BL treatments than either single mutant in the dark. (A) *Col-0*, *cca1-1*, *ataf2-2* and cca1-1 ataf2-2 showed similar hypocotyl lengths when grown in the dark. (B) *cca1-1 ataf2-2* did not show higher BL insensitivity than either *cca1-1* or *ataf2-2* in the dark. Four-day-old seedlings grown at 25 °C in darkness were used for hypocotyl measurements. Each data point represents the average of measurements from 30 seedlings (*n*=30). Error bars denote the standard deviations. n. s. *P*>0.01, \*\**P*<0.01, \*\*\**P*<0.001.

**Fig. S2.**
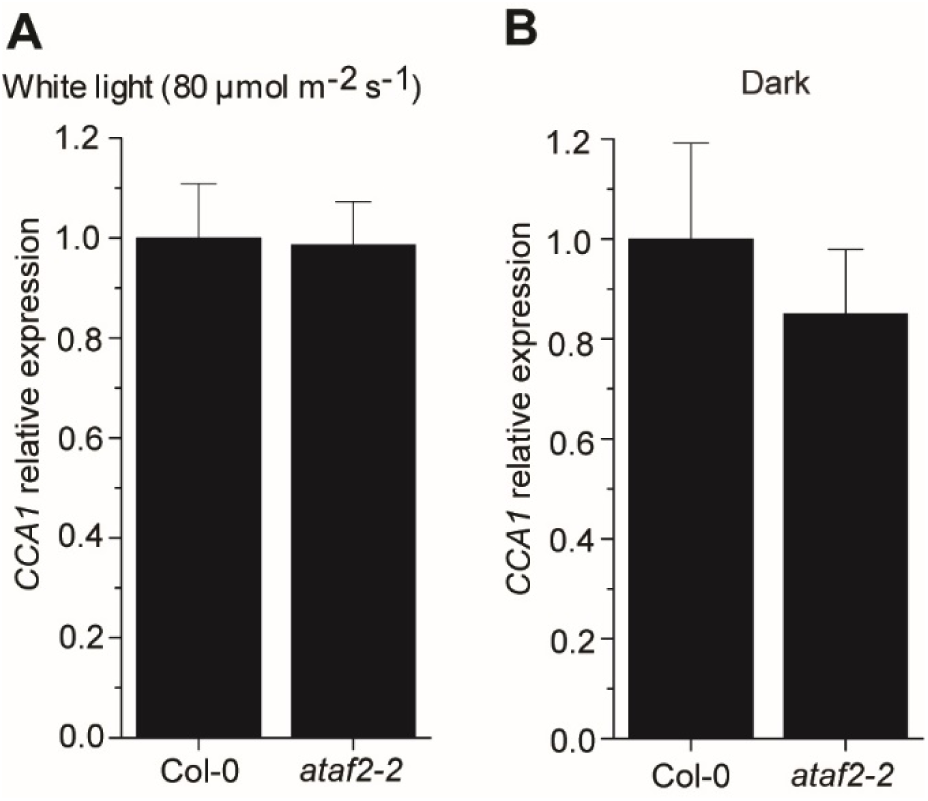
Compared to Col-0, *ataf2-2* seedlings did not show significant changes of *CCA1* expression in either continuous light (A) or darkness (B). Each qRT-PCR value is the mean of results from three biological replicates × three technical replicates (*n*=9). Error bars denote the s.e.m.

**Table S1.**
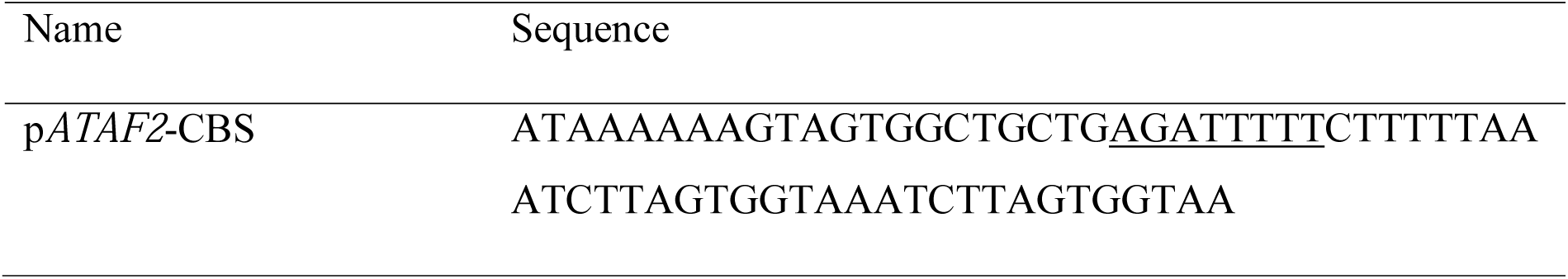
Sequences of *ATAF2* promoter fragment used for targeted Y1H. The CCA1 binding site (CBS) is underlined.

